# Filamin-A susceptibility to calpain-mediated cleavage as a marker of dynamic conformational changes in intact platelets

**DOI:** 10.1101/307397

**Authors:** Lorena Buitrago, Barry S. Coller

**Affiliations:** Allen and Frances Adler Laboratory of Blood and Vascular Biology, The Rockefeller University, 1230 York Ave New York, NY, USA 10065

**Keywords:** Calpain, Cytoskeleton, Filamin, Mechanotransduction, Platelet, integrin, platelet glycoprotein Ib

## Abstract

Filamin-A (FlnA), an actin-binding protein that organizes the actin cytoskeleton and mechanically links transmembrane glycoproteins to the cytoskeleton, associates with platelet receptors integrin αIIbβ3, glycoprotein-Ib (GPIb), and integrin α2β1. Fibrinogen, von Willebrand Factor (vWF) and collagen, binding to these receptors mechanically connect the extracellular matrix to the cytoskeleton. Here we identified that under standardized conditions, platelet activation and ligand binding to αIIbβ3, GPIb, or α2β1, generates reproducible patterns of FlnA cleavage after platelet lysis. We exploited this novel assay to study the impact of ligand binding and receptor activation on the platelet cytoskeleton. We identified that: i) the FlnA modification that renders it cleavable by calpain after platelet lysis, requires both ligand binding and either internal force (e.g; clot retraction) or external forces (e.g; stirring and aggregating), ii) FlnA modification depends on actin polymerization downstream of integrin αIIbβ3 and integrin α2β1, but not downstream of GPIb signaling, iii) FlnA modification is reversible in time when platelets are activated with convulxin (Cvx), collagen, von Willebrand factor (vWF) or thrombin receptor activating peptide (T6), and this reversibility correlates with platelet dissagregation, iv) in contrast to the reversible nature of platelet aggregation and FlnA modification by activation with Cvx, collagen, ristocetin or T6, when platelets are activated by thrombin the platelet do not disaggregate and FlnA remains cleavable. Our data demonstrate that αIIbβ3, α2β1 and GPIb can each exert tension on the cytoskeleton by virtue of binding ligand under conditions of shear. We further identified a unique role for αIIbβ3-fibrin interactions in creating sustained cytoskeletal tension, with implications for thrombus stability and clot retraction.

## Introduction

The platelet cytoskeleton plays a vital role in platelet physiology. Filamin A (FlnA; Actin-binding protein) is a major component of the cytoskeleton and one of the most abundant platelet proteins. Composed of two 280 kD subunits that dimerize at their C-termini, each subunit contains 24 immunoglobulin (Ig)-like domains organized in an N-terminal domain [comprised of an F-actin binding domain (ABD), followed by Ig 1-15] and a C-terminal domain [comprised of Ig 16-23 and a dimerization domain, Ig 24]. FlnA also contains two flexible hinge regions that contain calpain cleavage sites, H1 between Ig 15-16 and H2 between Ig 23-24 (Figure 1) (1, 2).

**Figure 1:**
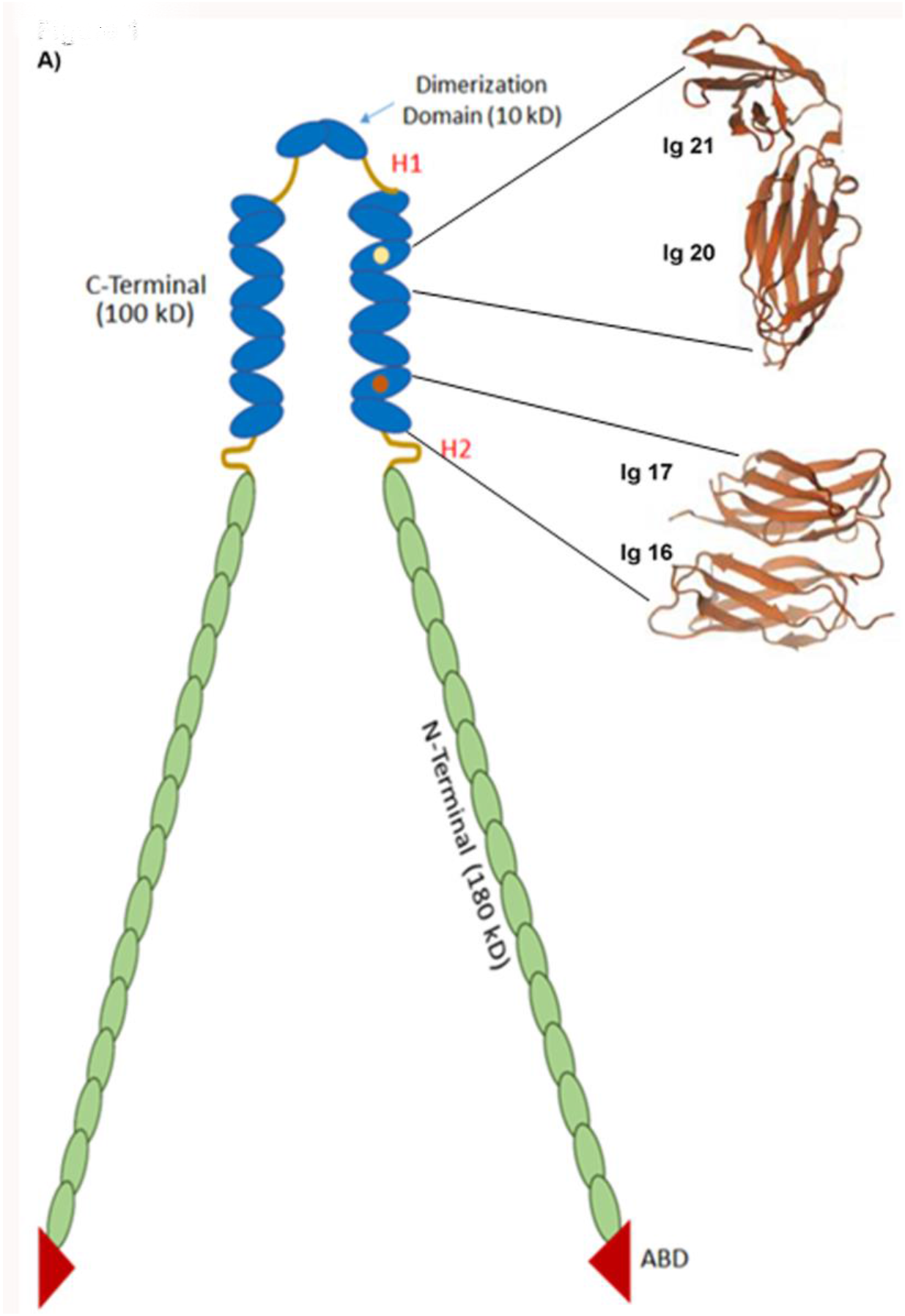
**A)** Schematic representation of Filamin A (FlnA). FlnA is a V-shaped dimer. Each monomer is 280 kD and contains 24 Ig-like repeats. FlnA can be divided into 2 domains: an N-terminal domain containing an F-Actin-binding domain (ABD) and repeats 1-15, and a C-terminal domain containing repeats 16-24. Domains 1617,18-19, 20-21 and 2-23 interact with each other. The C-terminal domain contains most of the partner interaction domains. Depicted are integrin αIIbβ3 interaction domain Ig 21 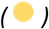 and GPIb interacting domain Igl7 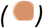. The C-Terminal also contains the dimerization domain Ig 24. FlnA hinge regions are located between Ig 23-24 (H1) and between Ig repeats 15-16 (H2). Hinge regions are susceptible to calpain cleavage. (Adapted from Dr. Mohammad R. K. Mofradd with permission. https://www.researchgate.net/publication/263853077_Filamin_A_Structural_and_Functional_Biomolecule_with_Important_Roles_in_Cell_Biology_Signaling_and_Mechanics).

FlnA organizes the cytoskeleton, stabilizes the plasma membrane, and contributes mechanical stability and rigidity to cells (3, 4). FlnA physically links membrane glycoproteins to the cytoskeleton by dynamically interacting with the cytoplasmic domains of the glycoproteins, F-actin, and several other cytoplasmic binding partners, including kinases and transcription factors, primarily through binding sites on its C-terminal domain (5, 6). The major interaction sites within the C-terminal domain are in the odd-numbered Igs, specifically Igs 17, 19, and 21 (7). In this domain, the even-numbered Igs interact with odd-numbered Igs as pairs, resulting in variable occlusion of the binding sites for other proteins. These binding sites become exposed when tension causes conformational changes in FlnA that dissociate the odd-numbered Igs from their even-numbered partners (8–10)). In turn, the newly exposed Ig regions recruit interacting signaling molecules that convert the mechanical force into signaling events (4, 8, 11, 12).

The role of FlnA as a mechano-sensing protein implicated in cell migration, motility, membrane stability, and mechano-protection has been studied extensively (13–17). In platelets, FlnA has been implicated in contributing to several processes, including: megakaryopoiesis and platelet formation (18), supporting membrane integrity under high shear (17, 19), and serving as a negative regulator of integrin αIIbβ3 activation (20, 21). The GPIbα-FlnA interaction appears to be crucial for both normal megakaryopoiesis and platelet membrane stability since patients who lack GPIb (Bernard Soulier syndrome) have giant platelets that show decreased resistance to aspiration-induced deformation (22) and mice lacking either GPIb or FlnA have large platelets (23). Moreover, in murine models, the interaction between GPIb and FlnA has been shown to be indispensable for maintaining mechanical integrity of the platelet membrane during adhesion to vWF under conditions of flow (17). The linkage between GPIb and FlnA has been localized to a constitutive interaction involving Phe568/Trp570 in the GPIbα cytoplasmic tail and FlnA Ig 17 (6, 17, 24), although an interaction between the GPIbα cytoplasmic tail with Ig21 has also been reported (8).

The integrin αIIbβ3-FlnA interaction is less well defined than the GPIb-FlnA interaction and the physiological implications are less well characterized. Several studies have demonstrated an interaction between integrin β7 subunit tail peptides (which have limited homology to the corresponding regions of β3) and FlnA Ig 21 (8, 25, 26). Liu *et al* (20) directly studied the interaction of FlnA Ig 21 with αIIb and β3 cytoplasmic domain peptides and found that Ig 21 has two binding sites for β3 and another binding site for αIIb. Thus, the tripartite complex between Ig 21, αIIb and β3, may potentially maintain the association of the cytoplasmic domains and presumably, therefore, inhibit αIIbβ3 activation. Alternatively, or additionally, FlnA has been proposed as a negative regulator of integrin activation based on the overlap between the β3 integrin subunit binding domain for FlnA with those for talin and kindlin, since the binding of these two proteins is required for receptor activation. According to this model, FlnA has to be displaced from the integrin tail, perhaps by migfilin (27), in order for talin and kindlin to bind and activate the integrin (25, 27). It is unclear, however, whether αIIbβ3 interacts with FlnA under basal conditions since the Ig 21 binding sites are occluded by its interaction with its partner Ig 20.

Most of data described above have come from structural studies on isolated proteins and peptides rather than studies of intact cells over time. Thus, one of the remaining goals of FlnA biology is to define the dynamics of FlnA conformational changes in cells as a function of cell activation and exposure to both external and internal sources of tension. In the course of our studies of the role of FlnA in platelet function, we discovered a novel method for assessing a modification in FlnA structure in intact platelets, namely the susceptibility of FlnA to calpain cleavage after platelet lysis under tightly controlled conditions. This report describes the impact of agonist-specific ligand binding to αIIbβ3, GPIb, GPVI, and α2β1 in conjunction with either internal or external forces in inducing either reversible or irreversible modifications in FlnA. Our results have implications for understanding cytoskeletal mechano-transduction in platelets as well as thrombus stability and clot retraction under shear.

## Materials and Methods

All reagents were obtained from Sigma-Aldrich unless stated otherwise. Anti-FlnA monoclonal antibody (mAb) clone TI10 (C-terminal) and Anti-FInA mAb CLB228 (N-Terminal) were purchased from EMD Millipore Corp. Mouse mAb clone TA205 to the talin head domain was purchased from Upstate Cell Signaling solutions. mAbs to GPIb (6D1) and to α2β1 (6F1) were produced by our laboratory (28, 29). The αIIbβ3 antagonist RUC-4 was prepared in our laboratory (30). A protease and phosphatase inhibitor cocktail (containing aprotinin, bestatin, E64, leupeptin, sodium fluoride, sodium pyrophosphate, β-glycerophosphate, and sodium orthovanadate) and β-actin rabbit mAb 13E5 purchased from Cell Signaling Technologies. Platelet agonists included: a thrombin receptor activating peptide (T6, SFLLRN) synthetized by the Rockefeller University Proteomics Resource Center; collagen Type I from rat tail purchased from Corning Inc; Convulxin (Cvx) a generous gift from Dr. Satya Kunapuli (Temple University); ristocetin obtained from Sigma-Aldrich, von Willebrand factor and human α-thrombin purchased from Haematologic Technologies Incorporated, and 2-methylthio-ADP trisodium salt (2-MeADP) purchased from Santa Cruz Biotechnology. All reagents were analytical grade.

### Studies with Human Platelets

This study was approved by the Rockefeller University Institutional Review Board (IRB). Human blood was obtained from healthy volunteers with informed consent who had not taken anti-platelet medications during the 2 weeks before blood collection and anti-coagulated with one-sixth volume of acid-citrate-dextrose (2.5 g of sodium citrate, 1.5 g of citric acid, and 2.0 g of glucose in 100 ml of H_2_O). Platelet-rich plasma (PRP) was prepared by centrifugation of citrated blood at 650 × *g* for 4 min at room temperature. To prepare washed platelets, prostaglandin E1 was added to ACD-PRP to a final concentration of 1 μM, and the preparation was centrifuged at 980 × *g* for 8 min at room temperature. The platelet pellet was resuspended in HEPES modified Tyrode’s buffer (HBMT) (138 mM NaCl, 2.7 mM KCl, 2 mM MgCl2, 0.42 mM NaH2PO4, 5 mM glucose, 10 mM HEPES, pH 7.4). The suspended platelets were allowed to regain their reactivity by incubation at 37°C for 45 min. For platelet aggregation studies, the washed platelet count was adjusted to 2 × 10^8^ platelets/mL and for studies of clot retraction the count was adjusted to 3 × 10^8^ platelets/mL.

### Platelet aggregation

In vitro platelet aggregation was evaluated at 37°C in a photometric aggregometer (Platelet Aggregation Profiler PAP8; Bio Data Corporation). Aggregation was induced by adding the indicated agonist and stopped by adding an equal volume of cold Tris-Triton lysis buffer (2% v/v Triton X-100, 100 mM Tris-HCl, pH 7.4) or one-third volume 3X SDS sample buffer [250 mM Tris-HCl (pH 6.8), 8% (w/v) sodium dodecyl sulfate (SDS), 0.2% (w/v) bromophenol blue, 40% (v/v) glycerol, 20% (v/v) β-mercaptoethanol]. When platelets were lysed with Tris-Triton buffer, the lysed samples were incubated with one-third volume of SDS sample buffer, and heated to 100°C for 10 min before analyzing by SDS polyacrylamide gel electrophoresis and immunoblotting. In some experiments, platelet lysis was performed with Tris-Triton lysis buffer with the added protease inhibitors, as indicated.

### Fibrinogen binding

To assess αIIbβ3 fibrinogen binding, washed platelets at a concentration of 2× 10^6^ platelets/mL were activated with the indicated agonists for 30 min at 37 °C under static conditions in the presence of 100 μg/mL Alexa647-conjugated fibrinogen. Platelets were then washed and resuspended in HBMT buffer and analyzed by flow cytometry (FACS Calibur).

### Immunoblotting

Washed platelets were stimulated with agonists at 37°C in an aggregometer cuvette for the indicated times, and the reaction was stopped by the addition of one-third volume SDS sample buffer or ice-cold Tris-Triton Buffer. In the latter case, total cell lysate was incubated for 1 hr on ice, and then one-third volume of SDS sample buffer was added. 50 μl samples were electrophoresed in 6-10% SDS-polyacrylamide gels. Proteins were electro-transferred to a nitrocellulose membrane (Millipore) and the membrane was then blocked for 1 hr with a proprietary blocking buffer (LI-COR) and incubated for 16 hr with the appropriate antibody 1:1000 (v/v) in the blocking buffer with 0.1% (v/v) Tween 20 at 4 °C. After washing, membranes were incubated for 1 hr at room temperature with species-specific, infrared-labeled secondary antibodies in a 1:20,000 dilution (LI-COR). Protein bands were visualized using an infrared detector (LI-COR, Odyssey). To directly assess the equivalence of protein loads, an anti-actin antibody was used for immuno-detection of actin content.

### Clot retraction

A suspension of 300 μL of washed platelets at a concentration of 3 × 10^8^ platelets/mL was added to a glass cuvette, mixed with 1 mM CaCl2 and either 0.2 or 2 units/mL of thrombin. At the indicated time points, an equal volume of ice-cold Tris-Triton lysis buffer was added and samples were kept on ice for 1 hr, after which one-third of 3X SDS sample buffer was added. The lysate was further incubated at 37°C for 12 hr to fully dissolve the clot before analysis by immunoblotting.

### Statistical analysis

Results are expressed as means ± SD with the number of observations indicated as n. Data were analyzed using GraphPad Prism 6.07. Comparison between samples was determined using the Student *t* test. Differences were considered significant at *P* ≤ 0.05.

## Results

### Platelet activation modifies FInA, rendering it cleavable by calpain after platelet lysis

In accord with studies by Fox et al. (31, 32), who used thrombin to activate platelets, we first observed that when washed platelets aggregated for 2 min in response to T6 and were lysed with a Tris/Triton lysis buffer and analyzed by SDS-PAGE under reducing conditions, that there was time-dependent FlnA cleavage. Cleavage of full-length FlnA (280 kD) produced three distinct fragments: an N-terminal domain (180 kD) observed with a FlnA N-terminal-specific mAb (Figure 2A, left panel), and both a full-length C-terminal domain (100 kD) and a cleaved C-terminal domain (90 kD) observed with a C-terminal-specific mAb (Figure 2A, right panel). When studied as a function of time, FlnA cleavage correlated with the onset of platelet aggregation, and the appearance of the 100 kD and 90 kD FlnA C-terminal fragments correlated with the potency of the aggregation response (Figure 2B, left panel). In sharp contrast to these results, when platelet samples were lysed directly with SDS sample buffer, the cleaved fragments were barely detectable (Figure 2B, right panel). These data suggested that while the cleavage of FlnA depended on activation by T6, the cleavage was actually occurring after platelets lysis. To test this hypothesis, we added to the Tris/Triton-X lysis buffer a commercial protease inhibitor cocktail (containing a proprietary mixture of leupeptin, aprotinin, bestatin, and E64), the cysteine protease inhibitor leupeptin, and the calpain-specific inhibitor calpeptin. The protease inhibitor cocktail had little or no effect on FlnA cleavage, whereas the higher concentrations of leupeptin and calpeptin partially inhibited the cleavage (Figure 2C). In contrast, adding the potent inhibitor of sulfhydryl proteases N-ethylmaleimide (NEM) at 20 mM to our Tris/Triton-X lysis buffer nearly completely inhibited FlnA cleavage induced by T6 (Figure 2D). To assess whether other platelet agonists could also lead to FlnA cleavage before or after platelets lysis, we induced platelet aggregation by activating platelets with the P2Y12 and P2Y1 agonist 2-MeADP, the GPVI agonist Cvx, the GPVI and α2β1 ligand collagen, or the inducer of von Willebrand factor (vWF) binding to GPIb, ristocetin, in the presence of added vWF. As with T6, all of these agonists induced FlnA cleavage when the aggregated samples were lysed without calpain inhibitors, but addition of 50 nM of calpeptin partially inhibited cleavage, and addition of 20 mM NEM completely inhibited the cleavage of FlnA (Figure 2D). Notably, the majority of FlnA molecules underwent cleavage with all of the agonists if maximum aggregation was achieved. We concluded from these results that upon platelet activation and aggregation FlnA is modified in a way that renders it susceptible to cleavage by calpain after platelet lysis. Moreover, since both the 100 kD and 90 kD C-terminal fragments were produced, it suggested that the conformation of FlnA is changed in at least two different ways, one of which exposes the H2 cleaving site exclusively and the other in which both H1 and H2 cleavage sites are exposed.

**Figure 2:**
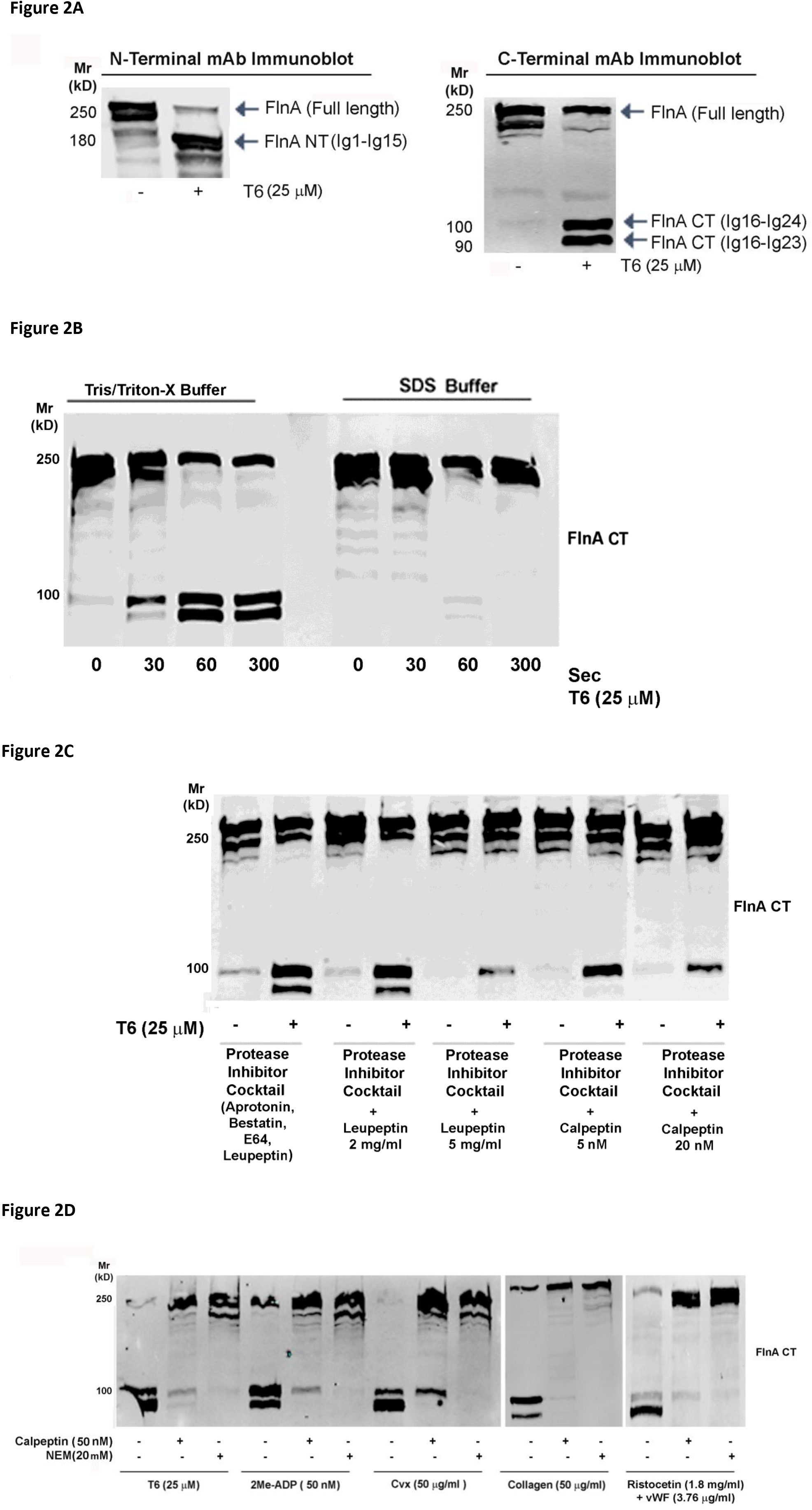
Platelet activation by T6, convulxin (Cvx), 2Me-ADP, collagen and ristocetin all lead to cleavage of FlnA after lysis and the cleavage can be prevented by inhibitors of the sulfhydryl protease calpain. **A)** Washed platelets were activated with 25 μM thrombin receptor activating peptide (T6) and reaction stopped after 2 minutes by adding cold Tris/Triton-X buffer. Samples were subjected to SDS-PAGE under reducing conditions and probed with an FlnA N-terminal-specific antibody that recognizes full length (280 kD) and the cleaved N-terminal fragment (180 kD) (Left panel) or with a C-terminal-specific antibody that recognizes full length (280 kD) and cleaved forms (100 kD and 90 kD) of FlnA (Right panel). **B)** Platelets were activated with 25 μM of T6 and reactions were stopped at the indicated time points by adding cold Tris/Triton-X lysis buffer or one-third volume SDS sample buffer. Samples were subjected to SDS-PAGE under reducing conditions and probed with FlnA C-terminal-specific antibody. **C)** Platelets were activated with 25 μM of T6 and the reaction stopped after 2 minutes by adding of cold Tris/Triton-X buffer containing the indicated protease inhibitors. Samples were subjected to SDS-PAGE under reducing conditions, and immunoblotted with an FlnA C-terminal antibody, **D)** Platelets were activated with the indicated agonists, and the reaction stopped after 2 min by adding cold Tris/Triton-X lysis buffer containing high concentrations of the calpain inhibitor calpeptin or the sulfhydryl protease inhibitor N-ethylmaleimide (NEM). Blots are representative of 3 experiments. FlnA-CT=FlnA C-terminal. FlnA NT=Filamin A N-Terminal

### FlnA modification requires ligand binding to integrin αIIbβ3, GPIb or α2β1

We next assessed whether ligand binding is required for platelet agonists to produce the modification in FlnA that renders it cleavable by calpain. Treating platelets with the αIIbβ3 antagonist RUC-4 prevented FlnA modification when platelets were activated with 2-meADP and T6, but had no effect when platelets were activated by collagen or ristocetin + vWF (Figure 3A). FlnA modification produced by ristocetin-induced binding to vWF could be prevented, however, by treating platelets with a mAb to GPIb (6D1) (Figure 3B). We then assessed the potential roles of integrin αIIbβ3, α2β1 and GPVI in the FlnA modification produced by collagen. We used Cvx to selectively activate platelets through GPVI (33) and inhibited collagen binding to α2β1 by treating platelets with the monoclonal antibody 6F1 (29). As seen in Figure 3C, blocking α2β1 with mAb 6F1 did not inhibited the FlnA modification induced by Cvx, but it partially inhibited the collagen-induced modification. Cvx-induced FlnA modification was, however, partially inhibited by RUC-4, but not by 6F1, suggesting that the Cvx-mediated FlnA modification is mediated in part by ligand binding to αIIbβ3. Taken together our results indicate that FlnA modification requires ligand binding to integrin αIIbβ3, GPIb, or α2β1. This was further supported by finding that PMA induced Ca^++^ mobilization and inside-out signaling but failed to induce the modification in FlnA if ligand binding to αIIbβ3was inhibited by RUC-4 or ligand binding to GPIb was inhibited by mAb 6D1 (Supplemental Figure 1). Thus, inside-out signaling in absence of ligand binding is insufficient to induce the modification in FlnA.

**Figure 3.**
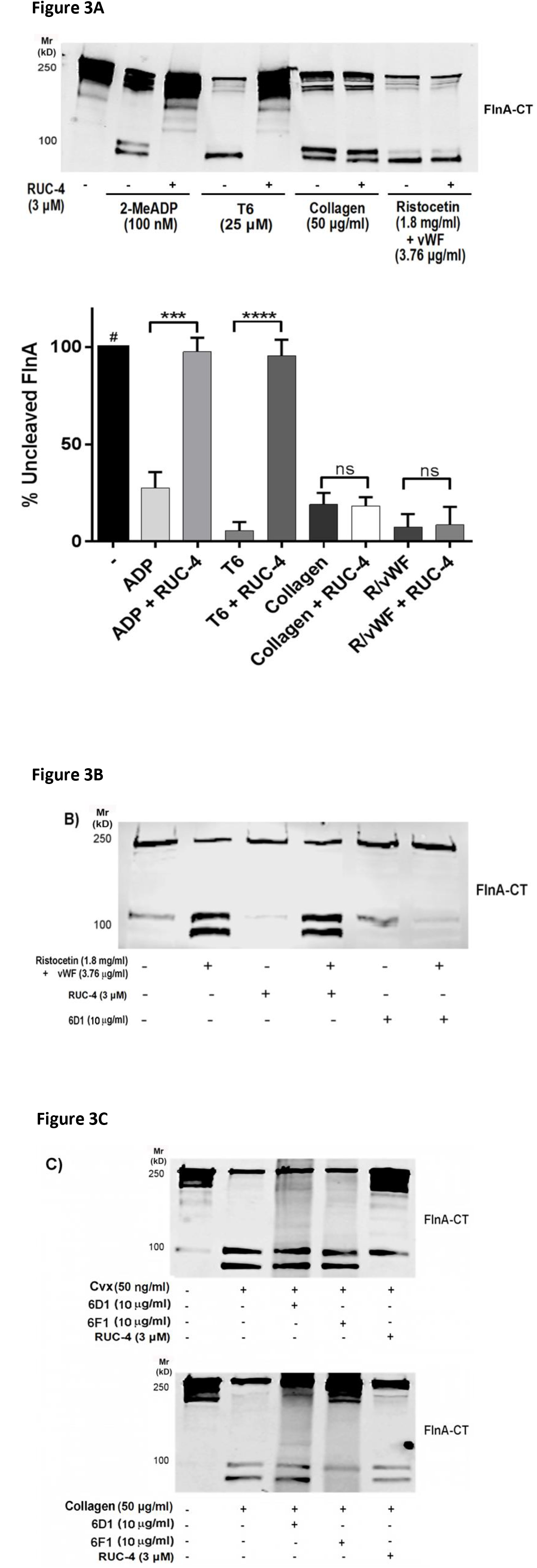
Ligand binding is required for platelets activation to result in the modification of FlnA that renders it susceptible to calpain cleavage. **A)** Effect of RUC-4 on FnA cleavage induced by 2-MeADP, T6, ristocetin + vWF and collagen. Washed platelets were activated with the indicated agonist and the reaction was stopped after 2 minutes by the addition of cold Tris-Triton lysis buffer. Some platelet preparations were treated with the integrin αIIbβ3 inhibitor RUC-4 at 3 μM for 20 minutes prior to activation. Top: Lysates were subjected to SDS-PAGE and immunoblotted with FlnA C-Terminal antibody. Bottom: Densitometry analysis of 4 experiments. Bars represent mean ± SD of full length FlnA (280 kD). **** = p<0.0001 ***= p< 0.001. **B)** Effects of RUC-4 and anti-GPIb mAb 6D1 on FlnA cleavage induced by ristocetin (1.8 mg/ml) + vWF (3.76 μg/ml) for 2 minutes. **C)** Effect of mAb 6D1, mAb 6F1, and RUC-4 on modification of FlnA induced by Cvx (top panel) or collagen (lower panel). Washed platelets were treated with Cvx (50 ng/ml) or collagen (50 μg/ml) for 5 minutes. Blots are representative of 3 experiments.

### FlnA modification requires either external or cell-generated force

To assess whether external or cell-generated mechanical forces play a role in the FlnA modification, we compared FlnA cleavage in samples from stirred and unstirred platelets (external forces) and under conditions of clot retraction (cell-generated forces). As seen in Figure 4A, FlnA modification induced by T6, Cvx, 2-MeADP, collagen, or ristocetin + vWF occurred only under stirring conditions even though fibrinogen binding occurred with each of these agonists in the absence of stirring (Figure 4B). In contrast, the FlnA modification occurred with platelets activated with thrombin, even in the absence of stirring. This stands in contrast to activation induced by T6, which also activates the thrombin PAR-1 receptor, and suggests that thrombin-mediated fibrin formation provides a matrix on which the platelets can exert tension. To test this hypothesis, we induced clot retraction with thrombin in the absence of stirring and observed FlnA cleavage after just 5 min of clot retraction, a time point at which fibrin polymerization has occurred but the clot has not grossly retracted (Figure 4C). As in samples from aggregating platelets, addition of 20 mM NEM to the lysis buffer inhibited the calpain-mediated cleavage, indicating that cleavage occurred after lysis.

**Figure 4:**
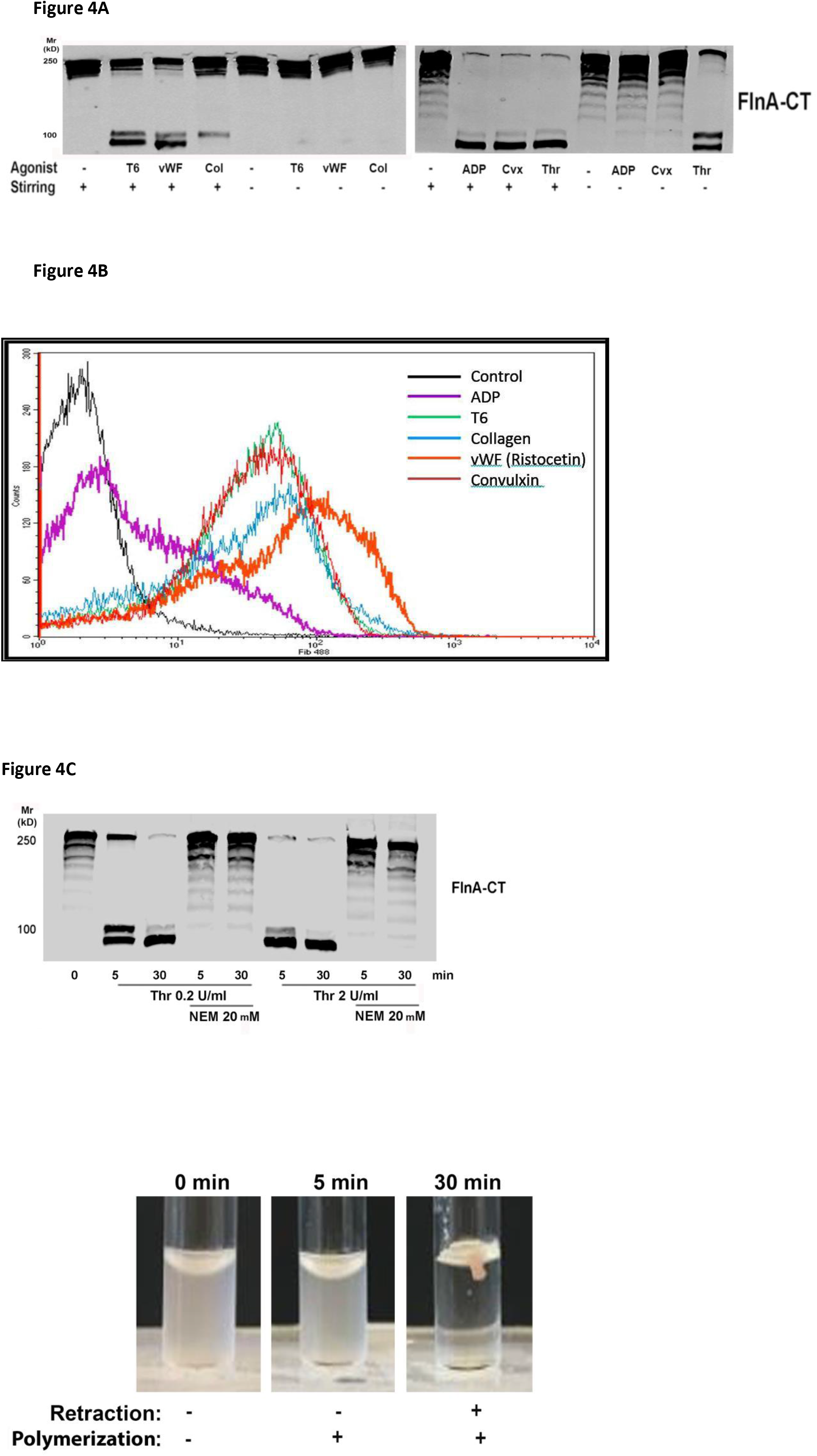
FlnA modification by T6 (25 μM), 2-meADP (100 nM), Cvx (50 ng/ml), or ristocetin + vWF (ristocetin 1.8 mg/ml and vWF3.76 mg/ml) occurs with stirring whereas stirring is not required when thrombin (0.2 U/ml) is used as an agonist or to initiate clot retraction. **A)** Washed platelets were treated with the indicated agonists for 5 minutes with or without stirring. Reaction was stopped by addition of cold Tris-Triton lysis buffer and samples processed for immunoblotting. **B)** Ligand binding occurs under static conditions. Platelets (2 x 10^6^ plat/ml) were activated with T6 (25 μM), 2Me-ADP (100 nM), Cvx (50 ng/ml), thrombin (Thr, 0.2 U/ml) or ristocetin + vWF (ristocetin 1.8 mg/ml and vWF 3.76 mg/ml) under static conditions without stirring, incubated with Alexa 647-labeled fibrinogen and analyzed by flow cytometry. The flow cytometry tracing is representative of 3 separate experiments. **C)** FlnA is modified during clot retraction. Washed platelets were treated with 0.2 or 2 U/ml thrombin and 1 mM CaCl*2* to initiate clot formation. At the indicated time points the reaction was stopped by adding cold Tris/Triton lysis buffer with or without 20 μM NEM and incubated for 1 hr. One-third volume SDS sample buffer was then added and the sample was further incubated at 37 °C for 12 hr to dissolve the clot. Bottom: Representative image of the retraction process at 5 and 30 min.

### FlnA modification requires actin polymerization downstream of ligand binding to integrin αIIbβ3 and integrin α2β1 but not downstream of GPIb

To assess a potential contribution of polymerized actin (F-actin) to the activation-dependent FlnA modification, we incubated platelets for 20 min before activation with 100 μg/ml of cytochalasin D, a compound that binds to actin filaments and inhibits subunit association and dissociation (34). As seen in Figure 5, cytochalasin D significantly inhibited FlnA modification when platelets were activated with T6 and thrombin; it had a minor, but significant, effect when platelets were activated with collagen, and no effect when vWF binding was induced with ristocetin.

**Figure 5.**
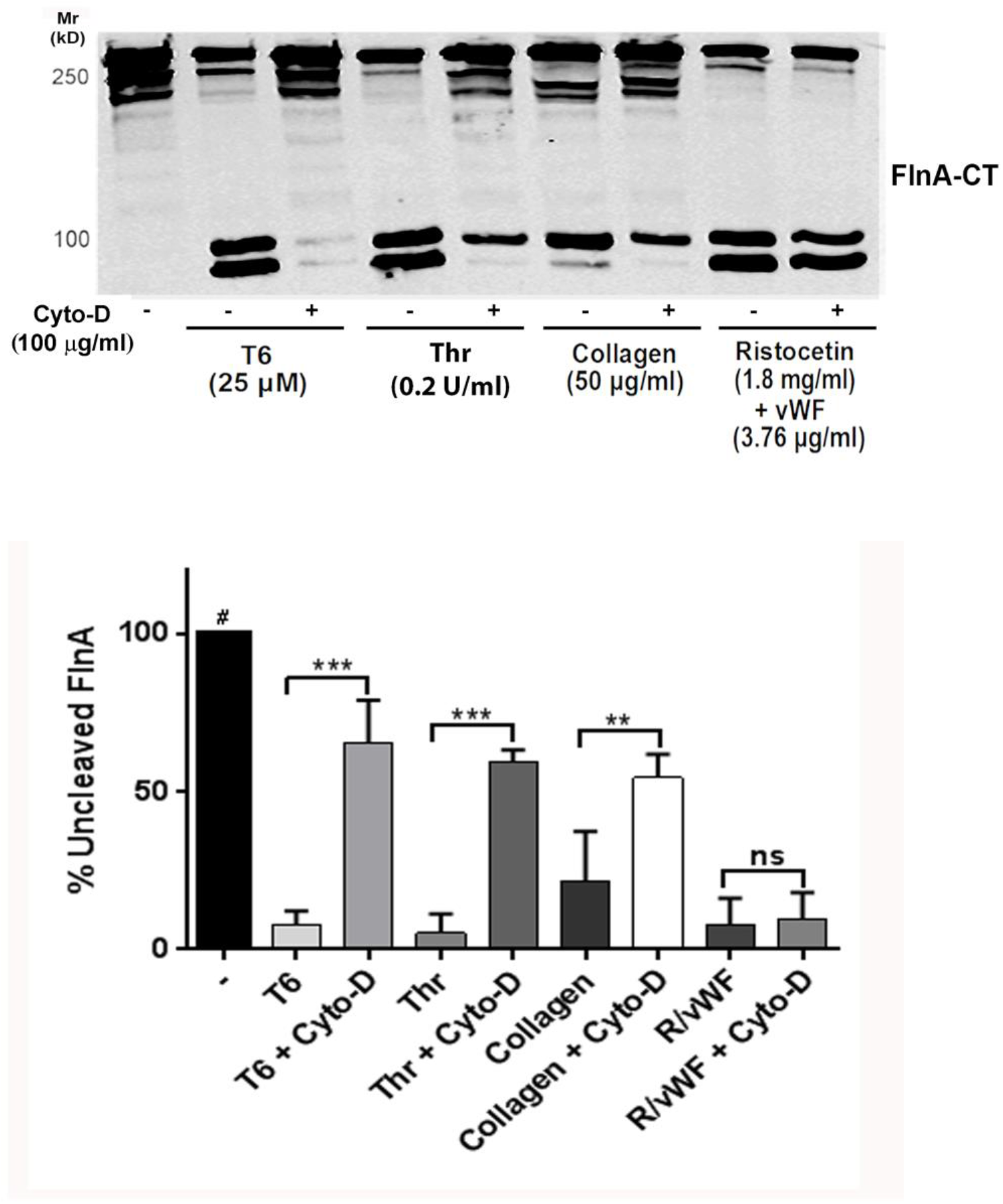
Cytochalasin D dissociates platelet activation from FlnA modification only platelets are activated with T6 or thrombin but not with ristocetin + vWF. Effects of cytochalasin D (Cyto-D) on FlnA cleavage induced by T6, thrombin (Thr), collagen and ristocetin + vWF (R/vWF). Washed platelets were activated with the indicated agonist and the reaction was stopped after 2 minutes by the addition of cold Tris-Triton lysis buffer. Some platelet preparations were treated with cytochalasin D at 100 μg/ml 20 minutes prior to activation. Top: Lysates were subjected to SDS-PAGE and immunoblotted with FlnA C-terminal antibody. Bottom: Densitometry analysis of 3 experiments. Bars represent mean ± SD of full length FlnA (280 kD). *** = p<0.001 **= p< 0.01

## FlnA modification downstream of ligand binding to αIIbβ3, GPIb, and α2β1 are reversible in time, but those downstream of fibrin binding to αIIbβ3 are not

To further explore the time course of the activation-dependent FlnA modification, we correlated the platelet aggregation and FlnA cleavage responses to the agonists over an extended time period. We observed that: 1) by 10 minutes after aggregation was initiated with T6, Cvx, collagen, or ristocetin + vWF, there was robust platelet aggregation and full-length FlnA (280 kD) was no longer detectable, indicating that nearly all of the platelet FlnA had undergone modification. 2) by 15-60 minutes after aggregation was initiated, aggregates began to disaggregate and some full-length, uncleaved FlnA was again identified by immunoblotting, indicating that the conformational change was at least partially reversible and that it correlated with the extent of platelet aggregation/disaggregation (Figure 6A). With T6, nearly quantitative cleavage of both H1 and H2 occurred in the 30 second and 1 minute samples after platelet lysis (resulting in the dominance of the 90 kD C-terminal fragment), whereas at 5, 10 and 30 minutes only about half of the molecules that underwent cleavage at H1 also underwent cleavage at H2. With collagen as the agonist, adding RUC-4 reduced the time until disaggregation occurred, and this correlated with immunoblot data demonstrating the reversal of the FlnA modification as judged by reduced FlnA cleavage at those time points (Figure 6B). We observed less disaggregation when ristocetin + vWF was used to activate the platelets and this reduced the reversibility of the FlnA modification. Adding RUC-4 facilitated the reversibility of aggregation and FlnA modification at 30 and 60 minutes (Figure 6C). In contrast to the reversible effect of the agonists described above, activating platelets with thrombin (0.2 U/ml) and stirring for as long as 60 minutes did not result in partial disaggregation or reversal of the conformational change in FlnA (Figure 6D). These data indicate that fibrin polymerization and binding to αIIbβ3 leads to irreversible platelet aggregation that correlates with an irreversible or prolonged change in FlnA conformation.

**Figure 6.**
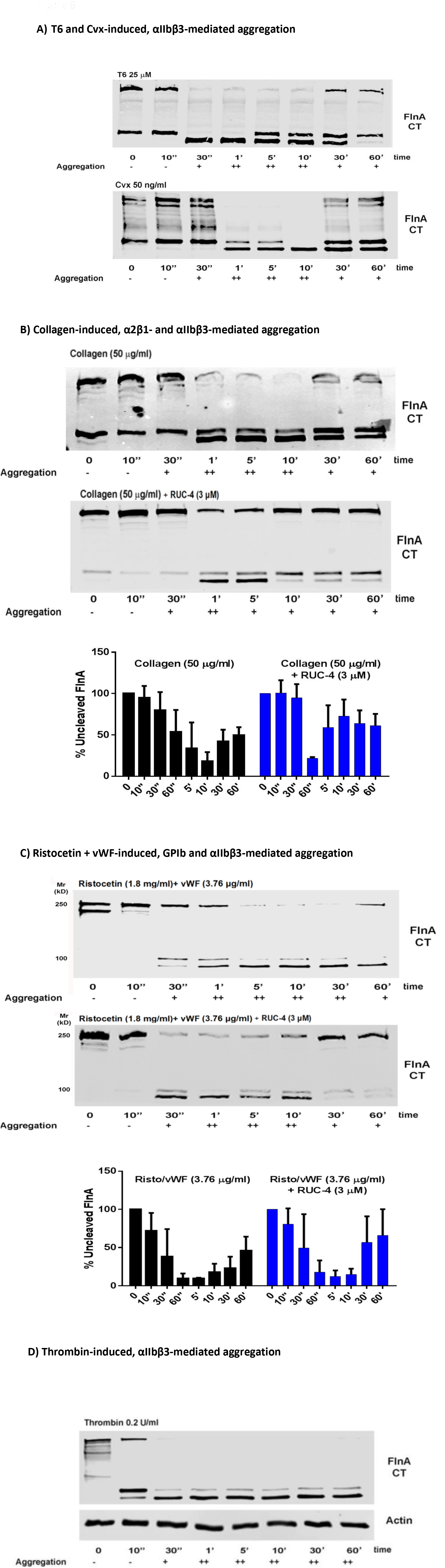
Time course of platelet aggregation and FlnA cleavage. Washed platelets (2×10^8^ platelets/ml) were activated under stirring conditions with the indicated agonist. At the indicated time point cold Tris/Triton lysis buffer was added and after 1 hr on ice, samples were analyzed by immunoblotting using a mAb that detects the C-terminal fragment of FlnA (FlnA CT). **A)** T6 and Cvx-induced aggregation, **B)** Collagen-induced aggregation in the absence or presence of RUC-4 inhibit ligand binding to αIIbβ3, lower panel contains densitometry analysis of 3 experiments. Bars represent mean ± SD of the full-length FlnA band (280 kD). **C)** Ristocetin/vWF induced aggregation with and without RUC-4, lower panel contains densitometry analysis of 3 experiments. Bars represent mean ± SD of the full-length FlnA band (280 kD). Note that the uncleaved FlnA band (Mw 280 kD) disappears or is significantly reduced after 5 minutes of aggregation but reappears at 10-30 min when disaggregation begins. In contrast, **D)** with thrombin activation, full-length FlnA disappears within 10 minutes but does not reappear and platelet disaggregation does not occur. Actin is used as a loading control. Platelet aggregation was judged to be + when the initial slope was between 10% and 50% and ++ when the initial slope was greater than 50%.

## Discussion

Although elegant structural studies of fragments of FlnA and its binding partners have defined interaction sites and the dynamics of the individual interactions under different experimental conditions, a full understanding of FlnA’s role in connecting the actin cytoskeleton to the platelet membrane glycoproteins and the impact of the associations on both inside-out and outside-in signaling requires understanding on how those interactions occur in intact platelets under both basal conditions and with activation, and under both physiologic and pathologic shear forces. In this study, we exploited our novel finding that platelet activation associated with ligand binding, in combination with internal or external forces, generates reproducible patterns of FlnA cleavage after platelet lysis under standardized conditions. The cleavage was inhibited by inhibitors of calpain, suggesting that the cleavage reflected conformational changes in FlnA that expose the H1 and/or H2 hinge regions, or perhaps less likely, conformational changes in molecules that ordinarily protect the hinge regions from cleavage. Thus, we were able to study the impact of ligand binding to GPIb, αIIbβ3, GPVI, and α2β1 in the presence and absence of external shear force induced by stirring or in the case of αIIbβ3, internal tension generated by clot retraction. Below we summarize our results by receptor in the context of current understanding of the molecular interactions.

### GPIb

There is strong evidence that GPIb is constitutively linked to FlnA via its cytoplasmic domain interacting with FlnA Ig 17 (6, 17, 24, 35), which does not appear to be auto-inhibited by Ig 18, and an additional interaction with Ig 21 has been proposed (9), although it is presumably subject to autoinhibition by the interaction with Ig 20. One or both of the interactions is vital for platelet membrane integrity and may contribute to GPIb-mediated shear-induced platelet aggregation (17). Consistent with these data, defects in GPIb in Bernard-Soulier syndrome in humans and mice result in large platelets, thrombocytopenia, and diminished membrane stability (36, 37) Genetic defects in the X-linked FlnA gene in humans result in a wide variety of clinical syndromes, some of which are associated with thrombocytopenia and large platelets, but normal or increased amounts of GPIb (38, 39). Platelets in these patients have been reported to demonstrate variable functional abnormalities (40). Targeted deletion of the FlnA gene in mice is embryonically lethal due to hemorrhage, cardiac defects, and abnormal vascular patterning, but conditional targeted deletion in erythroid cells, megakaryocytes and platelets results in female heterozygous mice with thrombocytopenia, modestly enlarged platelets, and long bleeding times, and male hemizygous mice with more severe thrombocytopenia and larger platelets. In both cases there is a decrease in GPIb (41).

Our data further support a functional interaction between GPIb and FlnA since we found that vWF binding to GPIb in the presence of stirring and platelet aggregation rapidly results in a conformational change in nearly all of the platelet FlnA, rendering it susceptible to calpain cleavage after platelet lysis. This modification is reversible over time and the reversal correlates with reversal of the extent of platelet aggregation. Some of the effects of vWF engagement of GPIb appear to be mediated by activation of αIIbβ3 as judged by the ability of the αIIbβ3 inhibitor RUC-4 to facilitate the reversal of both aggregation and the FlnA modification. This is of interest in light of the studies in mice with platelet FlnA deficiency showing the importance of Syk interaction with FlnA Ig 5 in αIIbβ3 activation by GPVI and CLEC-2 (41). Of particular note, unlike the conformational changes in FlnA produced by engaging integrin receptors, those produced by vWF binding to GPIb are not inhibited by disrupting actin polymerization with cytochalasin D. This is consistent with the binding of GPIb to FlnA being constitutive and the lack of auto-inhibition of Ig 17 (42).

### GPVI and α2β1

The interaction of Cvx with GPVI and collagen with GPVI and α2β1 also leads to the reversible conformational change in FlnA when platelets were stirred. Inhibiting ligand binding to GPIb did not affect either result, whereas inhibiting ligand binding to αIIbβ3 partially inhibited the Cvx-induced conformational change, but had little effect on the initial collagen-induced conformational change; it did, however, speed and enhance the reversal of the conformational change induced by collagen. Thus, signaling to FlnA through GPVI and α2β1 is variably effected through activation and ligand binding to αIIbβ3. Furthermore, since cytochalasin D decreased FlnA cleavage, actin polymerization also appears to contribute to the process.

### αIIbβ3

The interaction of αIIbβ3 with FlnA is less well defined than the interaction with GPIb. Based on studies of isolated Ig 21 in the absence of its auto-inhibiting binding partner Ig 20, FlnA has been proposed to inhibit inside-out signaling (20, 25, 43) by: 1) preventing the binding of talin, and perhaps kindlin, to the β3 cytoplasmic domain (25), and possibly 2) forming a ternary complex with both αIIb and β3 that prevents them from dissociating (20). It has further been proposed than migfilin plays an important role in integrin αIIbβ3 inside-out signaling and activation by displacing FlnA from β3 and allowing talin and kindlin free access the β3 cytoplasmic tail (27). These data still need to be reconciled however, with the observation that FlnA null platelets do not demonstrate constitutively active αIIbβ3 receptors (41) and the very small amount of migfilin present in platelets relative to FlnA. In particular, while the copy number for FlnA in human platelets has been estimated to be 87,700 molecules per platelet, migfilin was not on the list of platelet proteins in the proteomic analyses of either human or mouse platelets (44), and in unpublished studies we have found minimal amounts of migfilin by immunoblotting. Together, these data suggest that in contrast to GPIb-FlnA interactions, αIIbβ3-FlnA interactions are not constitutive; rather they require mechanical force to relieve the auto-inhibition due to the binding of Ig 20 to Ig 21, which occludes the binding site on Ig 21 for the β3 cytoplasmic tail (45).

Our data demonstrate that ligand binding to αIIbβ3 induced directly by T6 or ADP, or indirectly after ligand binding to GPIb, GPVI, or α2β1, leads to FlnA modification, but only when external force is applied via stirring. Even with activation by PMA, with documented intracellular signaling as judged by phosphorylation of target proteins, the induction of the conformational change in FlnA required both ligand binding and stirring. Furthermore, in contrast to the lack of effect of cytochalasin D on FlnA modification downstream of ligand binding to GPIb, cytochalasin D significantly decreased FlnA cleavage downstream of ligand binding to αIIbβ3, suggesting that polymerized actin plays an important role in the conformational change in FlnA, perhaps by transmitting the force required to dissociate Ig 20 from Ig 21, thus relieving the auto-inhibition and facilitating the interactions between αIIbβ3 and FlnA responsible for the conformational change. As with ligand binding to GPIb, the conformational change in FlnA is reversible over time as platelet aggregates decrease in size.

Activation of αIIbβ3 by thrombin differs from activation induced by the other agonists in that the conformational change occurred in the absence of stirring. It did, however, correlate with clot retraction, indicating that in this case, the tension required to induce the conformational change came from within the platelet as a result of interaction of actin and myosin. Moreover, the conformational change induced by thrombin was unique in that it was not even partially reversible within the 60-minute time period studied. This may be relevant to in vivo thrombus formation because, as platelet thrombi form on damaged blood vessels, the hydrodynamic drag produced by the flowing blood on the platelets in the thrombus increases and so the bond-strength required to prevent embolization increases. The generation of thrombin and the formation of fibrin are therefore important in stabilizing the thrombi since fibrin interacts differently with αIIbβ3 than fibrinogen (46), with the bond strength greatest with polymerized fibrin, intermediate with fibrin monomer, and lowest with fibrinogen (47). The presence of ancillary binding sites on fibrin, separate from the γ-404-411 sequence, and complementary binding sites on αIIbβ3, may account for the greater bond strength (48, 49).

Separate from its potential roles in αIIbβ3 inside-out signaling, our data demonstrate that FlnA is modified downstream of ligand binding to αIIbβ3 and thus plays a role in outside-in signaling. We speculate that the observed FlnA modification is due to the transmission of tension sufficient to expose the H1 and H2 calpain cleavage sites, separate Ig 20 from Ig 21, thus exposing the binding sites for β integrin cytoplasmic tails on Ig 21, and concomitantly separating other Ig pairs, exposing additional binding sites for other partners important for cytoskeletal rearrangement and focal adhesion formation such as Rho, Rac, Cd42 and Syk (1, 50).

In other cell types talin, which also contains an actin-binding domain, has been implicated in regulating outside-in signaling since it also can connect the integrin β subunit to the cytoskeleton (51, 52). A mouse model described in studies by Petrich et al. suggests that in platelets, FlnA rather than talin regulates outside-in signaling (53). In their studies, they describe the phenotypes of platelets from two mouse strains, one with a β3 Y747A mutation that disrupts β3 interaction with talin and FlnA, and the other with a β3 L746A mutation that only disrupts β3-talin interactions. Both mutants have defects in αIIbβ3 activation, agonist-induced fibrinogen binding, and platelet aggregation as expected for disrupting talin binding to αIIbβ3. Importantly, platelets from the β3-L746A mice, but not the β3-Y747A mice, were able to support outside-in signaling as judged by their ability to spread on fibrinogen and phosphorylate FAK on tyrosine when αIIbβ3 ligand binding was induced by incubating platelets with MnCl2 in the presence of fibrinogen. Of particular note, while both animals had increased bleeding the severity was greater in the mice with the β3 Y747A mutation (5% of L746A mice vs. 53% of Y747A mice).

Data on the interactions among FlnA, αIIbβ3, GPIb, actin, and talin need to be considered in the context of the very high copy number of all of these molecules in platelets, with GPIb the lowest at ~25,000 copies per platelet. The elegant structural and single molecule studies that have identified the interactions of the GPIbα cytoplasmic domain with FlnA Ig 17 and Ig 21 (8, 17), and the interaction of FlnA Ig 21 with both membrane proximal and distal regions of the integrin β3 subunit as well as the αIIb subunit cytoplasmic domain, demonstrate that all of these interactions are likely to have short life-times, albeit probably longer for GPIb than β3 (based on studies of the related integrin β7). The combination of high copy number and short bond life-times allows for dynamic changes while still retaining constant connections at any time.

While calpain activation has been reported in association with platelet activation (54–56) the reversible nature of the FlnA conformational change we observed when platelets were activated with a variety of activators (other than thrombin) and the ability of potent calpain inhibitors to prevent FlnA cleavage when added to the lysis buffer (even with thrombin), support the hypothesis that calpain cleavage of FlnA occurs after lysis. Platelet lysis is expected to activate both the μ-calpain (which requires micromolar concentrations of Ca^++^ for activation) and m-calpain (which requires millimolar concentrations of Ca^++^ for activation) present in platelets since it will release Ca^++^ from intracellular storage sites (56). Evidence from mice lacking μ-calpain indicate that this enzyme does not contribute to cleavage of FlnA with platelet activation (57). There are no data currently available for mice lacking m-calpain because targeting the m-calpain gene is embryonically lethal. A role for μ-calpain in platelet aggregation has been proposed based on studies in μ-calpain^-/-^ mice, so there may be sufficient release of intracellular Ca^++^ with platelet activation to activate μ-calpain (but perhaps not m-calpain), but it appears that the reduced aggregation response in μ-calpain^-/-^ mice reflects the loss of inactivation of the phosphatase PTP1b rather than an effect on FlnA or other cytoskeletal proteins (58). Fox et al. previously observed FlnA cleavage with platelet activation by thrombin and inhibition of the cleavage by preventing ligand binding with an RGD peptide or mAb 10E5 (55). She concluded that the cleavage did not occur after platelet lysis because it was not inhibited by 0.1 mg/ml of leupeptin, but in our studies we found that even 2 mg/ml of leupeptin was unable to prevent the cleavage of FlnA after lysis. Thus, fully inhibiting calpain activation after lysis requires high concentrations of calpain inhibitors.

FlnA can be phosphorylated at many sites, and at some sites, phosphorylation may inhibit calpain-mediated cleavage (59), presumably by shielding the H1 and H2 hinge regions. It is possible, therefore, that release of phosphatases with platelet lysis may contribute to the enhanced FlnA cleavage we observed. However, our lysis buffers contained the phosphatase inhibitors sodium fluoride, sodium pyrophosphate, β-glycerophosphate, and sodium orthovanadate. In addition, we did not observe any change in the phosphorylation of residue S2152 (data not shown), which has been implicated in protecting FlnA from calpain cleavage with platelet activation or lysis (60, 61).

In conclusion, our data demonstrate that FlnA undergoes a modification upon platelet activation that renders it cleavable by calpain after platelet lysis. We used this novel system to study the platelet mechanical state and identified that αIIbβ3, α2β1 and GPIb can independently exert tension on the cytoskeleton by virtue of their binding ligand under conditions of shear. Our data also has potential implications for outside-in signaling and identifies a unique role for αIIbβ3-fibrin interactions in creating sustained cytoskeletal tension, with implications for thrombus stability and clot retraction.

## Authorship

Contribution: L.B. designed and performed research, analyzed data, and wrote the manuscript and B.S.C. designed research, analyzed data, and wrote the manuscript.

## Acknowledgments

This work was supported in part by the American Heart Association post-doctoral fellowship (award ID: 15POST25080127), the National Heart, Lung, and Blood Institute, National Institutes of Health (grant HL019278), and National Center for Advancing Translational Sciences, National Institutes of Health Clinical and Translational Science Awards (UL1TR001866), as well as funds from Stony Brook University. The authors thank the services of the High-Throughput and Spectroscopy Resource Center at Rockefeller University.

---

**Supplemental Figure 1:**
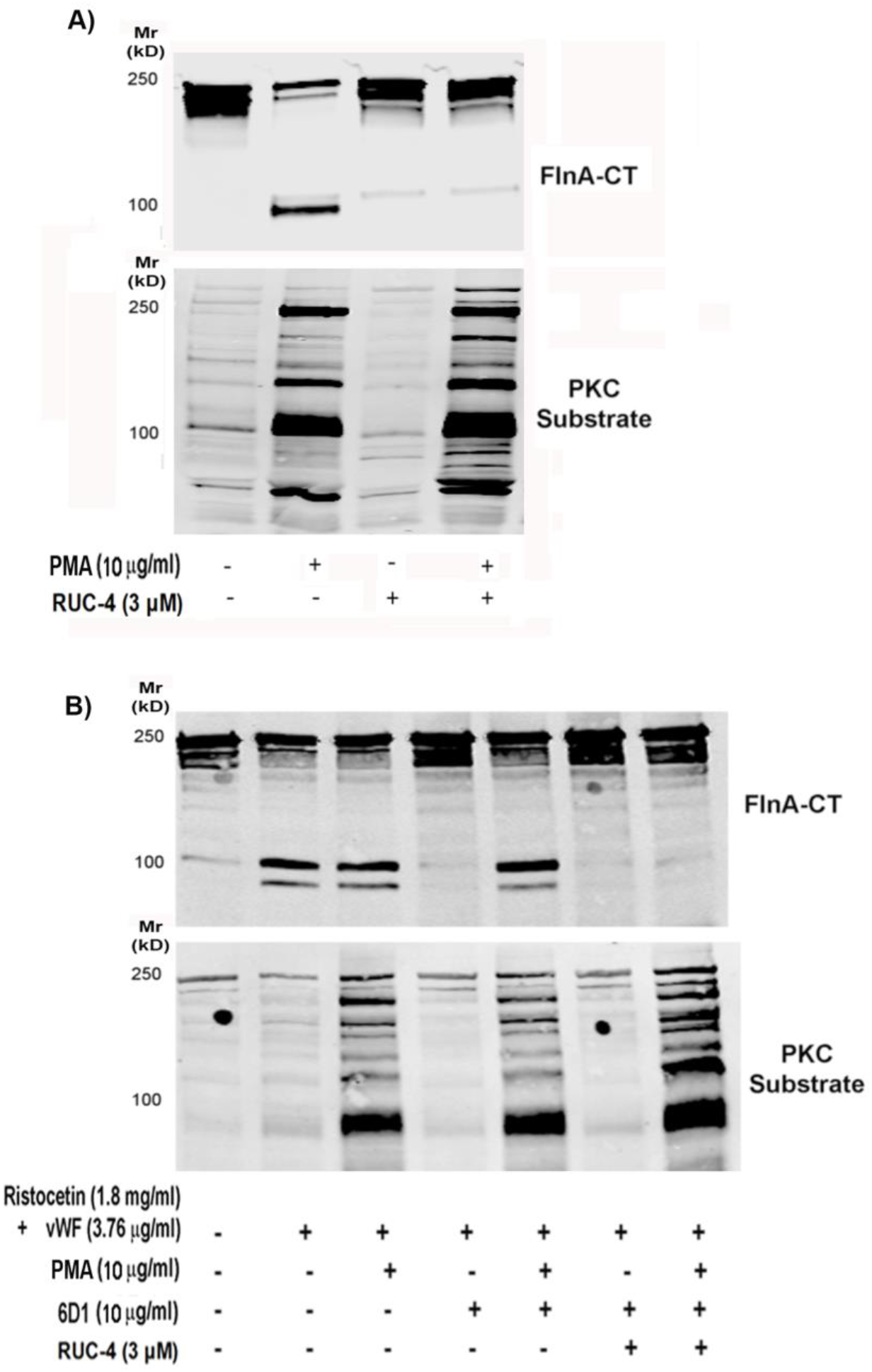
Inside-out signaling is not sufficient to induce FlnA modification. Washed platelets were treated with 10 μg/ml PMA for 10 minutes prior to be exposed to aggregating conditions. RUC-4 and 6D1 were added to inhibit ligand binding to αIIbβ3 and GPIb respectively. Immunoblot analysis with PKC substrate antibody demonstrate that PMA induces PKC activation, Ca++ mobilization and protein phosphorylation (inside-out signaling) but this is not sufficient to induce FlnA modification.

